# Abrogating ALIX interactions results in stuttering of the ESCRT machinery

**DOI:** 10.1101/576926

**Authors:** Shilpa Gupta, Mourad Bendjennat, Saveez Saffarian

## Abstract

ESCRTs are cellular proteins that catalyze the fission of membranes and play an important role in biology of disease including cancer and infectious virus release ^1^. ESCRT associated protein ALIX plays an essential role in HIV budding ^2-4^, exosome release ^5,6^, down regulation of G-protein coupled receptors ^7^, cytokinesis ^8,9^ and multi vesicular budding ^1^. The consensus view is that ALIX plays a role by binding to the viral late domains ^2-4^/Syntenin late domain ^5^/Cepp55 ^8,9^ and helps recruit downstream protein CHMP4 ^2,8,10,11^ which along with VPS4 catalyzes the fission of the membrane ^12,13^. Using live imaging we have visualized the recruitment of ALIX, CHMP4 and VPS4 during budding of HIV with abrogated Gag-ALIX interactions. Based on the canonical view, we were expecting to find reduced recruitment of ALIX under these conditions. Instead we report observing multiple rounds of transient recruitment of ALIX, CHMP4 and VPS4 prior to virion release. We further show that during each, transient recruitment, stoichiometry of all ESCRT components remained the same regardless of mutations abrogating ALIX and Gag interaction. In addition, mutations abrogating interactions between Gag and TSG101 result in recruitment of ESCRTs with a substantial delay while maintaining similar stoichiometry. Our results demonstrate that recruitment of ESCRTs is driven by a robust network of interactions resulting in an “On/Off” switch behavior and ALIX’s interactions with late domains of HIV Gag play a crucial role during final catalysis of membrane fission after assembly of the full ESCRT machinery.

---

We visualized the recruitment of ALIX during assembly of individual HIV Gag virus like particles (VLPs) on the basal membrane of HeLa cells stably expressing ALIX-h30-eGFP. ALIX-h30-eGFP links ALIX with eGFP through a stiff 30 amino acid super helical linker and is functional with the same efficiency as WT in rescue of PTAP^-^ HIV virion release ^14^. To generate fluorescent VLPs for fluorescent microscopy, cells were transfected with plasmids encoding HIV Gag-mCherry under a CMV promoter. Once VLP assembly commenced at the basal membrane, the membrane was imaged with a TIRF penetration depth of 150 nm using consecutive 488 nm and 561 nm illuminations every 15 seconds for 1.5 hours (methods). Individual VLPs were identified and analyzed from their initiation until full assembly which corresponds to a stable fluorescence signal from HIV Gag-mCherry as shown in Figure 1. From 40 VLPs analyzed we detected 75% single transient recruitment of ALIX at the end of assembly and 25% showing an average of two transient recruitment events.

**Figure 1:**
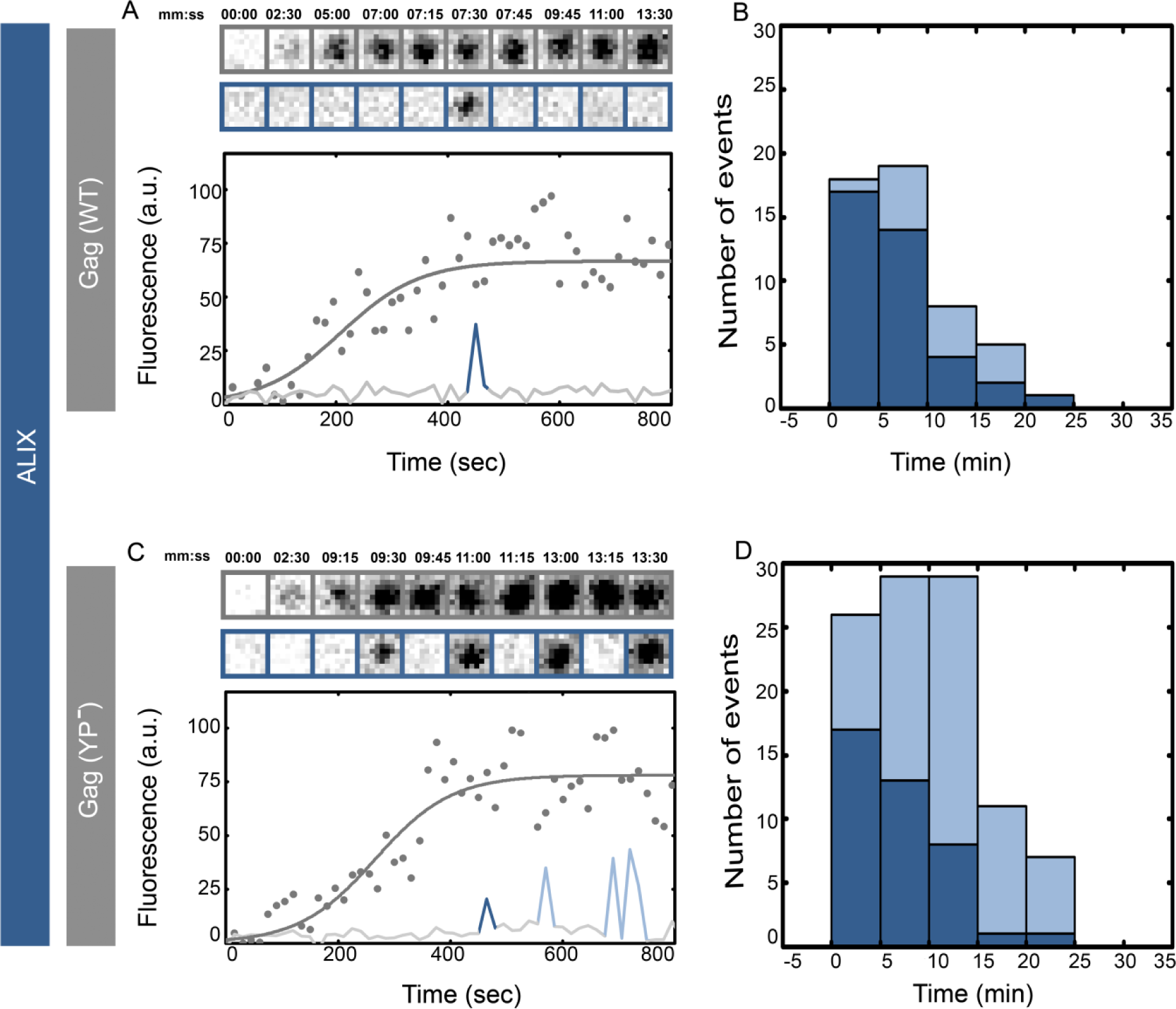
Single versus multiple transient recruitments of ALIX into HIV Gag VLPs versus HIV Gag(YP^-^) VLPs. HeLa cells stably expressing ALIX-h30-eGFP were transfected with 1500 ng of Gag-mCherry (A&B) and Gag(YP^-^)-mCherry (C&D) and imaged 4hours after transfection. Assembly of individual representative VLPs are shown with intensity plots and cropped TIRF images of the Gag (Top, Gray) and ALIX (bottom, Blue) for (A) Gag-mChery VLPs and (C) Gag(YP^-^)-mCherry VLPs. Histograms of the first time (dark blue) and later recruitments (light blue) of ALIX are shown for (B) Gag-mChery VLPs and (C) Gag(YP^-^)-mCherry VLPs. Majority of the first Alix recruitment is within 1-10 minutes after the VLP assembly completes during both YP^-^ and WT assembly.

ALIX interacts directly with HIV Gag through the YPXL late domain motif on p6 Gag ^2,7,15^. In Gag(YP^-^) we abrogated this interaction by incorporating (_36_SR_37_) in place of (_36_YP_37_) as previously characterized ^3^. We visualized the recruitment of ALIX during assembly of individual HIV Gag(YP^-^)-mCherry VLPs on the plasma membrane of cells stably expressing ALIX-h30-eGFP. Once VLP assembly commenced at the basal membrane, cells were imaged with identical settings to the imaging described above. The assembly times are not affected by the YP^-^ mutation as previously reported ^16-18^ and the average maximum intensity of HIV Gag(YP^-^)-mCherry VLPs were identical to wild type HIV Gag-mCherry VLPs (10500 ± 4000 a.u. versus 12000 ± 4000 a.u. respectively). After the completion of assembly however, YP^-^ VLPs transiently recruited multiple rounds of ALIX as shown in Figure 1. In 40 VLPs analyzed, 70% showed multiple rounds of recruitment ‘Stuttering’ with an average of 4 recruitment events per VLP. As shown in Figure 1 the arrival time of the first transient recruitment of ALIX in HIV Gag-mCherry VLPs was approximately 1-10 minutes post completion of assembly similar to the arrival of the first transient recruitment of ALIX into HIV Gag(YP^-^)-mCherry VLPs. The tagging of Gag with mCherry had no effect on assembly and the observed ALIX recruitment phenotype since similar results were obtained in experiments where VLPs assembled with a mixture of HIV Gag(YP-)-mCherry along with HIV Gag(YP-) as shown in Figure S1.

The intensity of the maximum fluorescent signal is proportional to the number of ALIX-h30-eGFP molecules recruited to the sites of virion release. Based on our analysis with the time resolution used in our study there was negligible difference between any transient ALIX recruitments into WT versus YP^-^ VLPs as shown in Figure S2. To verify that YP^-^ VLPs have released from the host cells and don’t remain tethered to the membrane we detached the cells from the glass using incubation in TryplE (Methods). Once the cells were removed the immobilized VLPs were visualized on the glass as shown in Figure S3.

ALIX has a well established biochemical interaction with CHMP4b through its Bro domain ^2-4,19^. We visualized the recruitment of CHMP4b during the assembly of individual VLPs on the plasma membrane. To visualize this recruitment during HIV Gag VLP assembly, we created a plasmid which expresses human CHMP4b linked to eGFP by a flexible linker at its N-terminus under a ΔCMV promoter (ΔCMV-eGFP-flex-CHMP4b). A similar N-terminally tagged CHMP4b has been used before to visualize recruitment of CHMP4b onto the assembling Gag VLPs by other labs ^20^. The co-expression of this plasmid had no effect on the release of HIV Gag VLPs as shown in Figure S4. In addition as shown in Figure S4, we further characterized this plasmid in infectious HIV release and found a slight decrease in virion release with no effect on infectivity of the released virions (Methods).

We visualized the recruitment of CHMP4b during assembly of HIV Gag VLPs on the plasma membrane of HeLa cells co transfected with ΔCMV-eGFP-flex-CHMP4b and HIV Gag-mCherry. Once VLP assembly commenced at the basal membrane, cells were imaged with identical settings to the imaging described above. Transient recruitment of CHMP4b was observed after completion of HIV Gag assembly as shown in Figure 2 consistent with previous observations of CHMP4b recruitment into assembling VLPs ^20^. From 40 VLPs analyzed, 80% showed a single CHMP4b recruitment and 20% showed an average of two CHMP4b transient recruitments.

**Figure 2:**
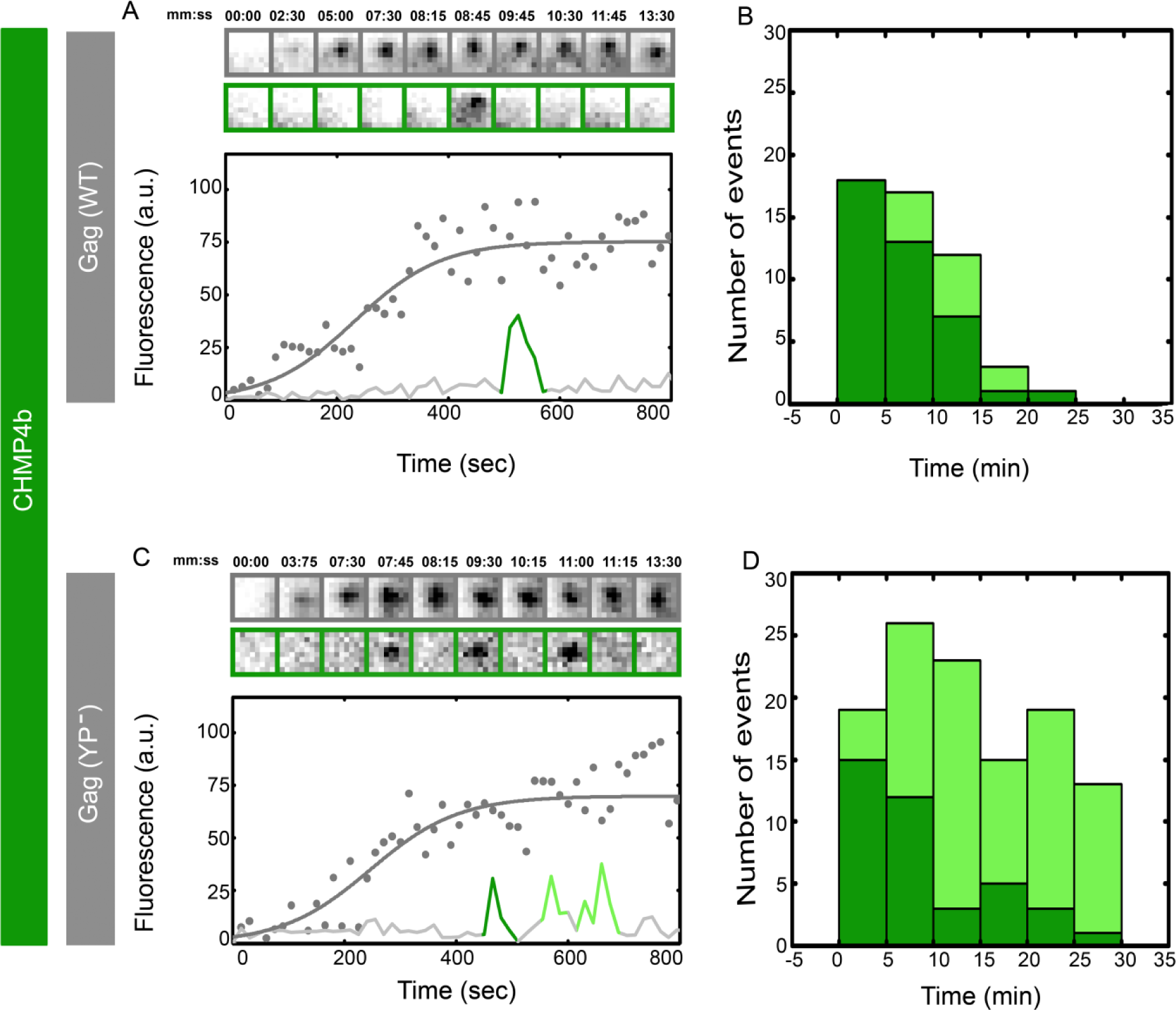
Single versus multiple transient recruitments of CHMP4 into HIV Gag VLPs versus HIV Gag(YP^-^) VLPs. HeLa cells were transfected with 900 ng of ΔCMV-eGFP-flex-CHMP4b and 600 ng of Gag-mCherry (A&B) and Gag(YP^-^)-mCherry (C&D) and imaged 7 hours after transfection. Assembly of individual representative VLPs are shown with intensity plots and cropped TIRF images of the Gag (Top, Gray) and CHMP4b (bottom, Green) for (A) Gag-mChery VLPs and (C) Gag(YP^-^)-mCherry VLPs. Histograms of the first time (dark Green) and later recruitments (light Green) of CHMP4b are shown for (B) Gag-mChery VLPs and (C) Gag(YP^-^)-mCherry VLPs. Majority of the first CHMP4b recruitment is within 1-10 minutes after the VLP assembly completes during both YP^-^ and WT assembly.

We further visualized the recruitment of CHMP4b during assembly of individual HIV Gag(YP^-^)-mCherry VLPs. As shown in Figure 2, CHMP4b was recruited transiently multiple times during assembly of HIV Gag (YP^-^) VLPs. From 40 VLPs analyzed, 75% showed multiple rounds of transient recruitment with an average of 4 recruitment events per VLP. The first transient recruitment of CHMP4b was 1-10 minutes after completion of HIV Gag assembly and was indistinguishable between Gag WT and YP^-^ VLPs. Similar to the ALIX recruitment, the peak intensity of CHMP4b during transient recruitments had a negligible difference between WT and YP^-^ VLPs as shown in Figure S5.

At the late stages of ESCRT function, CHMP4b and VPS4 are recruited to catalyze the fission of the membrane. We further visualized the recruitment of VPS4 into WT versus HIV Gag YP^-^ VLPs as shown in Figure 3. VPS4 proteins were linked to mCherry with a 37 amino acid long helical linker previously characterized^17^. HeLa cells were co-transfected with ΔCMV-VPS4-h37-mCherry along with HIV Gag-eGFP or HIV Gag(YP^-^)-eGFP and assembly of HIV Gag VLPs and recruitment of VPS4 were recorded and analyzed as described above. In 40 WT VLPs analyzed, 70% showed a single VPS4 recruitment and 30% showed an average of two transient recruitments. As shown in Figure 3, VPS4 is transiently recruited 1-10 minutes after completion of HIV Gag assembly with identical first recruitment timing between WT and YP^-^ VLPs. In 40 YP^-^ VLPs analyzed, 70% showed multiple rounds of VPS4 recruitment with an average of 3 recruitments per VLP. The peak intensity of VPS4 transient recruitments had a negligible difference between the WT versus YP^-^ condition as shown in Figure S6.

**Figure 3:**
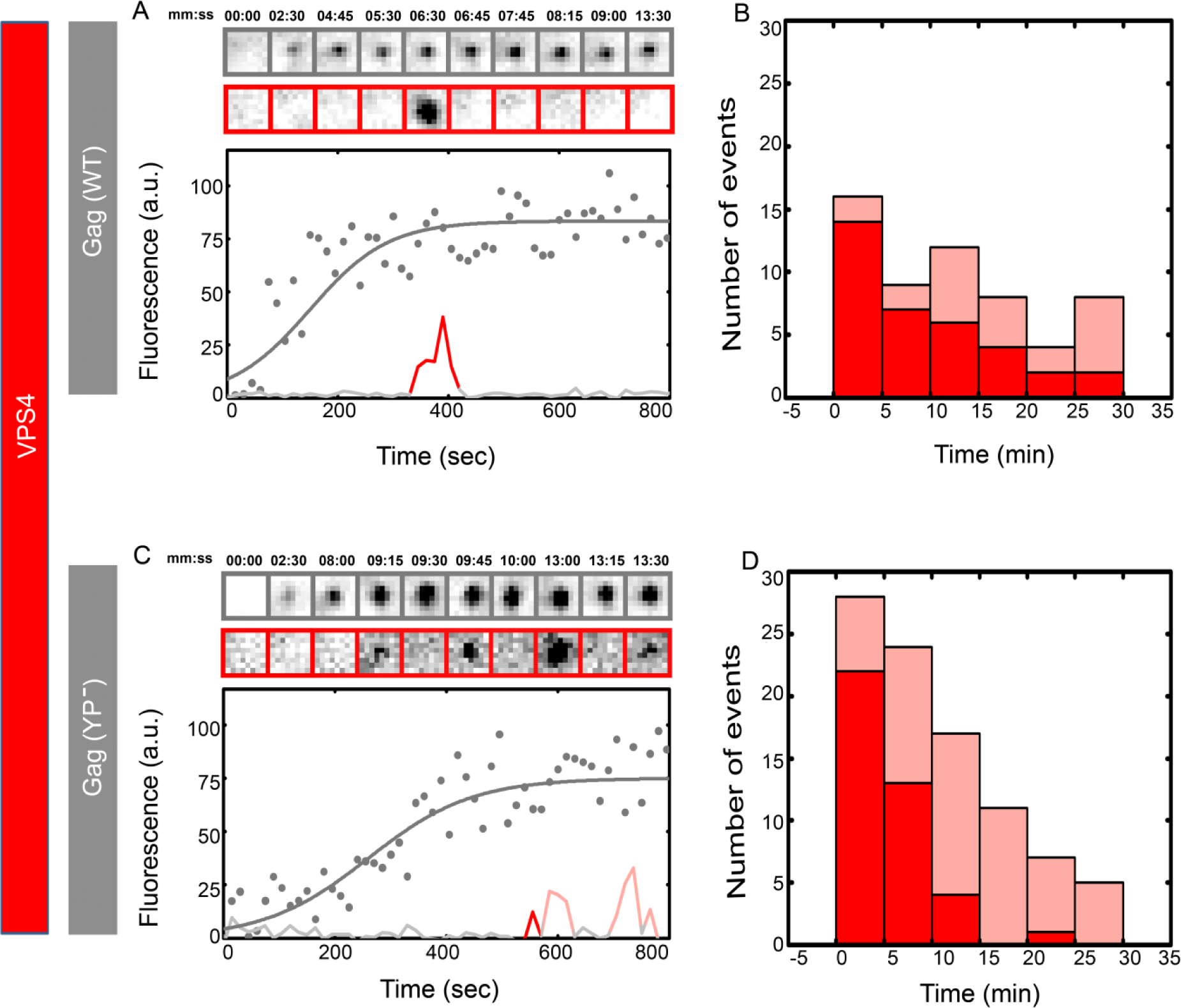
Single versus multiple transient recruitments of VPS4 into HIV Gag VLPs versus HIV Gag(YP^-^) VLPs. HeLa cells were transfected with 1200 ng of ΔCMV-VPS4-h37-mcherry and 300 ng of Gag-mCherry (A&B) and Gag(YP^-^)-mCherry (C&D) and imaged 7 hours after transfection. Assembly of individual representative VLPs are shown with intensity plots and cropped TIRF images of the Gag (Top, Gray) and VPS4 (bottom, Red) for (A) Gag-mChery VLPs and (C) Gag(YP^-^)-mCherry VLPs. Histograms of the first time (dark Red) and later recruitments (light Red) of VPS4 are shown for (B) Gag-mChery VLPs and (C) Gag(YP^-^)-mCherry VLPs. Majority of the first VPS4 recruitment is within 1-10 minutes after the VLP assembly completes during both YP^-^ and WT assembly.

Our experiments up to this point were carried out using HeLa cells and the HIV Gag protein. To verify if the observed stuttering of ESCRTs is more generally applicable and is not dependent on our particular experimental system we carried out the experiments in U2OS cells and also visualized the recruitment of VPS4 during assembly of NL4.3(iGFP)(ΔENV) VLPs. We visualized the assembly of HIV Gag VLPs in U2OS cells by transfecting ΔCMV-VPS4-h37-mCherry plasmid along with HIV Gag-eGFP or HIV Gag(YP^-^)-eGFP into these cells. While the assembly of both WT and YP^-^ VLPs in U2OS cells were slower, as shown in Figure S7 the recruitment of VPS4 showed similar behavior in U2OS cells as in HeLa cells. We also measured the recruitment of VPS4 onto individual HIV virions assembling on the plasma membrane of HeLa cells by co-transfecting ΔCMV-VPS4-h37-mCherry and NL4.3(iGFP)(ΔENV) or NL4.3(iGFP)(ΔENV)(YP^-^). As shown in Figure S8 the observed stuttering of VPS4 during assembly of NL4.3(iGFP)(ΔENV)(YP^-^) is similar to the stuttering observed in HIV Gag(YP^-^)-eGFP.

We further studied the effects of ALIX on the recruitment pattern of ESCRTs by depleting ALIX from cells using siRNA treatment. As shown in Figure S9 HeLa cells were depleted from ALIX after two rounds of the siRNA treatment as verified by the disappearance of the ALIX-h30-eGFP and accumulation of multinucleated cells due to ALIX depletion^9^. After two rounds of the siRNA, multinucleated HeLa-ALIX-h30-eGFP cells transfected by HIV-Gag-mCherry, assembled HIV Gag-mCherry VLPs did not show any ALIX recruitment as expected. However, under the same conditions, multinucleated HeLa cells transfected with ΔCMV-VPS4-h37-mCherry along with HIV-Gag-eGFP or HIV-Gag(YP^-^)-eGFP showed substantial stuttering of the recruitment of VPS4 as shown in Figure 4. The observed stuttering of VPS4 during budding of WT HIV Gag VLPs shows that the observed stuttering phenotype is present due to critical function of ALIX which is missing in both cells depleted of ALIX as well as in VLPs formed by the YP-mutated HIV Gag.

**Figure 4:**
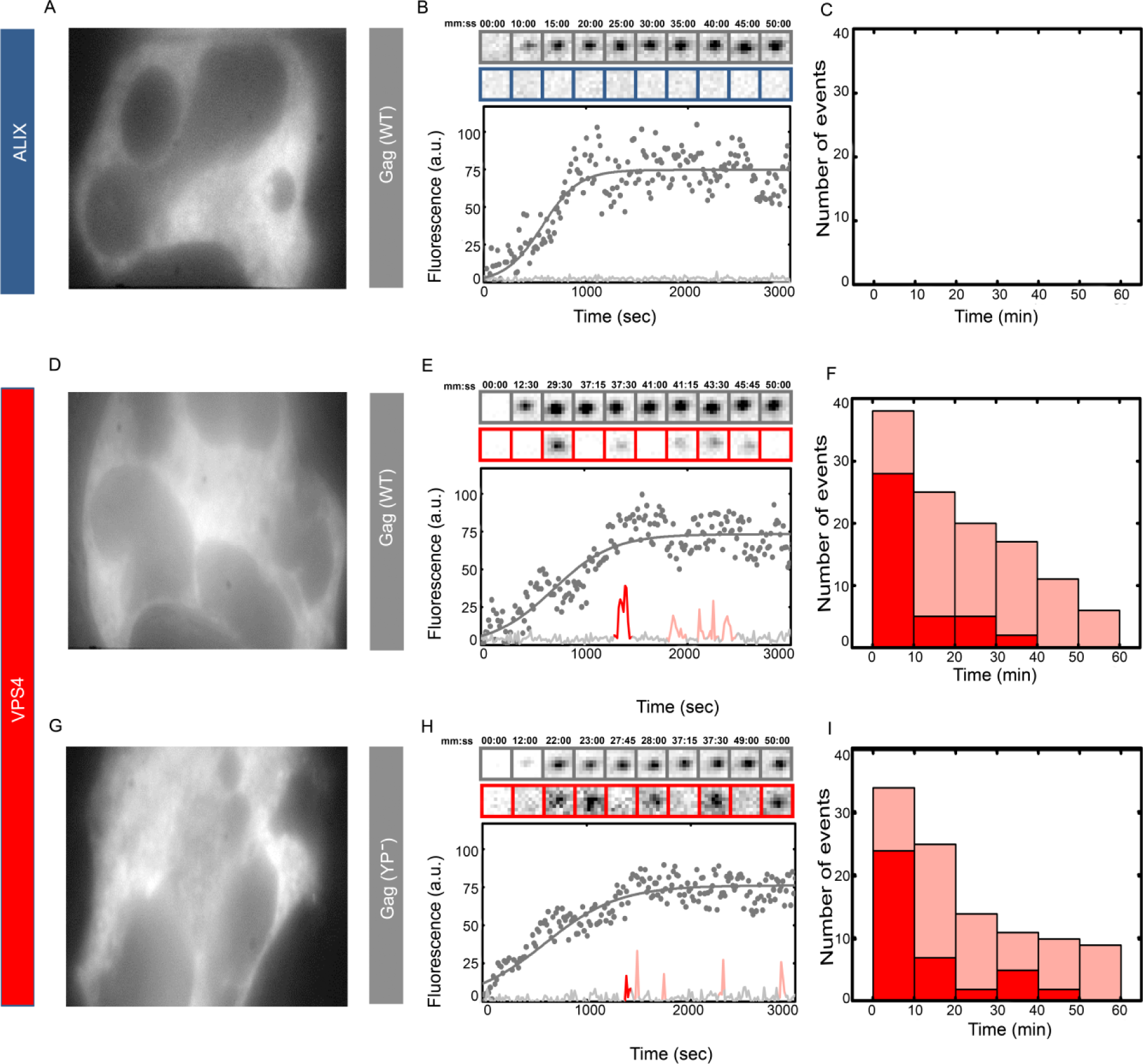
Multiple transient recruitments of VPS4 into HIV Gag WT and Gag(YP^-^) VLPs under ALIX depletion. Hela cells were treated with two rounds of siRNA against ALIX. (A, B & C) shows HeLa ALIX-h37-eGFP cell line treated with siRNA against ALIX and then transfected with 1500 ng of Gag-mCherry WT, an individual multinucleated cell was chosen (A) and a representative HIV Gag-mCherry VLP assembly in this cell is shown in (B) with intensity plots and cropped TIRF images of the Gag (Top, Gray) and ALIX (bottom, Blue). In 40 VLPs analyzed, there was no recruitment of ALIX as shown in (C). HeLa cells were treated with siRNA against ALIX and then transfected with 1200 ng of ΔCMV-VPS4-h37-mcherry and 300 ng of HIV Gag-eGFP (D, E & F) and Gag(YP^-^)-eGFP (G, H & I). (D&G) show multinucleated cells chosen for experiments with (E&H) showing a representative HIV Gag-mCherry (E) or HIV Gag(YP^-^)-mCherry (H) VLP assembly with intensity plots and cropped TIRF images of the Gag (Top, Gray) and VPS4 (bottom, Red). (F & I) Show histograms of the number and timing of the first (dark red) and later recruitment (light red) of VPS4 in (F) HIV Gag-mCherry VLPs and (I) HIV Gag(YP^-^)-mCherry VLPs.

There are two late domain motifs identified in the HIV Gag-p6. PTAP motif has been shown to directly interact with TSG101 and is critical for release of infectious HIV virions from infected cells ^21-23^. We further visualized the recruitment of ALIX into HIV Gag VLPs with both PTAP^-^ which incorporates a (_7_LIRL_10_) instead of (_7_PTAP_10_) ^24^ and PTAP^-^ + YP^-^ which has both PTAP and YP sequences altered (_7_LIRL_10_ plus _36_SR_37_). HeLa cells stably expressing ALIX-h30-eGFP were transfected with HIV Gag(PTAP^-^)-mCherry or HIV Gag(PTAP^-^+YP^-^)-mCherry and assembly kinetics and recruitment of ALIX were visualized with TIRF imaging as described above. During the initial experiments, conducted similarly to previously described experiments, no recruitment events were observed on the HIV Gag VLP assembly sites. To probe a later time frame recruitment, we therefore, initiated the TIRF imaging one hour after the initial formation of VLPs. In these sets of experiments, the VLPs are already fully assembled at the initiation of plasma membrane imaging and therefore exact timing of recruitment events to the completion of assembly time cannot be deduced. These experiments showed single transient recruitment of ALIX-h30-eGFP at a time frame between 1 hour and 2 hours post assembly of VLPs as shown in Figure 5. From 40 VLPs analyzed, 80% had a single recruitment event and 20% showed an average of two recruitment events. The multiple recruitments observed in the YP^-^ VLPs were not observed in the PTAP^-^ VLPs. The transient recruitment of ALIX was identical between PTAP^-^ and PTAP^-^ + YP^-^ mutation as shown in Figure 4. The intensity of the ALIX-h30-eGFP in transient recruitments had a negligible difference between WT, PTAP^-^ and PTAP^-^+YP^-^ VLPs as shown in Figure S10.

**Figure 5:**
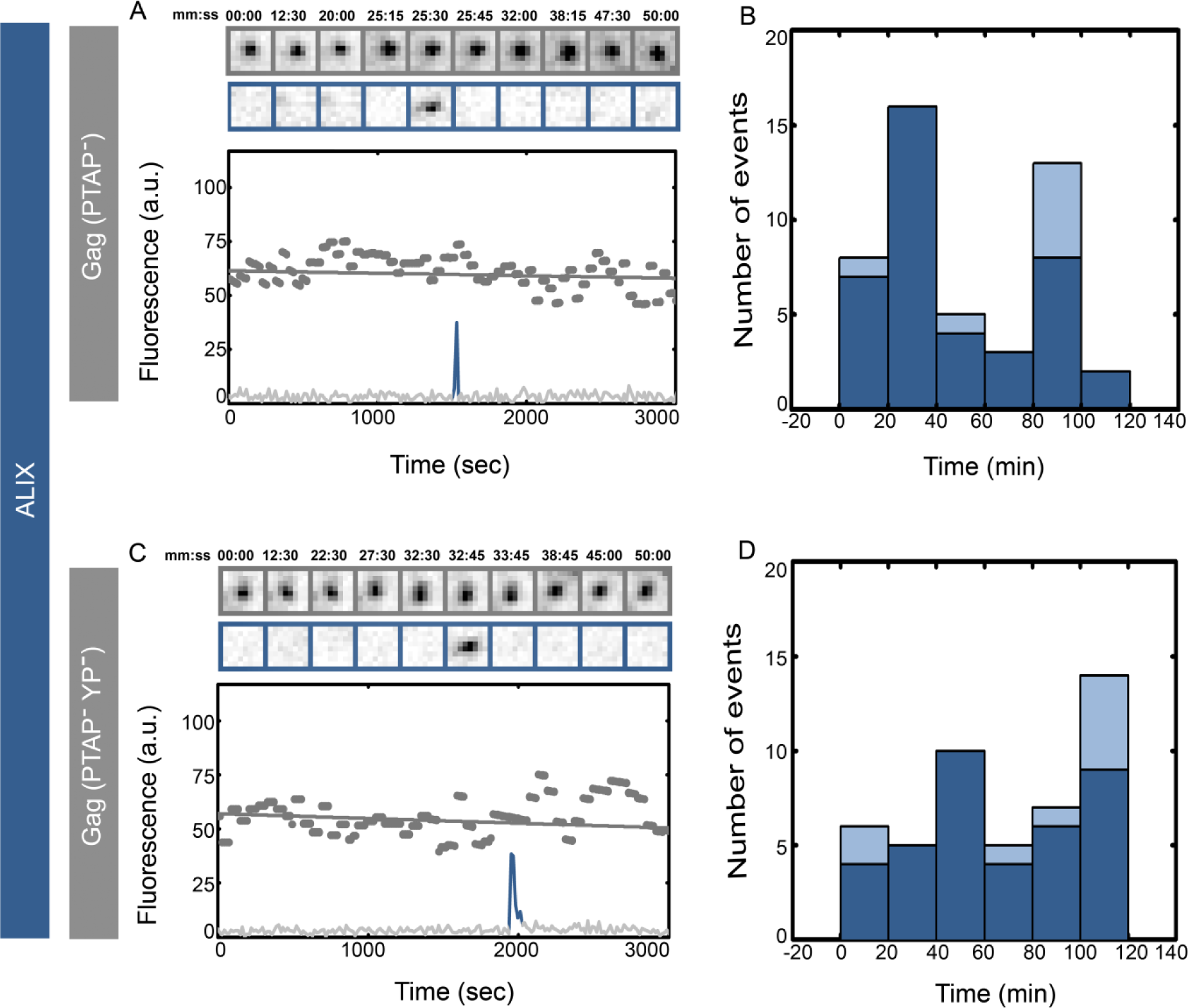
Single versus multiple transient recruitments of ALIX into HIV Gag(YP^-^ + PTAP^-^) VLPs versus HIV Gag(PTAP^-^) VLPs. HeLa cells stably expressing ALIX-h30-eGFP were transfected with 1500 ng of Gag(PTAP^-^)-mCherry (A&B) and Gag(PTAP^-^+YP^-^)-mCherry (C&D) and imaged 5 hours after transfection. Assembly of individual representative VLPs are shown with intensity plots and cropped TIRF images of the Gag (Top, Gray) and AIX (bottom, Blue) for (A) Gag(PTAP^-^)-mChery VLPs and (C) Gag(PTAP^-^+YP^-^)-mCherry VLPs. Histograms of the first time (dark blue) and later recruitments (light blue) of ALIX are shown for (B) Gag(PTAP^-^)-mChery VLPs and (C) Gag(PTAP^-^+YP^-^)-mCherry VLPs. Alix gets recruited within 2 hrs after the start of the actual experiment.

We suggest that virions are released during the last recruitment of ESCRTs in the stuttering transient recruitments. This suggestion is based on similarity of the measured delay in virion release, previously characterized in kinetic viral budding assays and our observed durations of the ESCRT stuttering on individual virions. In more detail, we have previously shown that mutations within the late domain of HIV result in a delayed release of the virus, which in turn results in budding of non-infectious virions due to premature protease activation ^25^. These kinetic biochemical assays showed an approximate delay of 20 minutes for HIV Gag (YP^-^) in U2OS cells and 70 minutes for fully infectious virions with YP^-^ mutation ^25^. This delay is consistent with the time between the first recruitment of ESCRTs and the last recruitment of ESCRTs on individual VLPs visualized in this study. Our measurements show an average time between these recruitments as 9.5 ± 7.5 minutes for HIV Gag (YP^-^) in HeLa cells, 18 ± 13 minutes for HIV Gag (YP^-^) in U2OS cells and 37 ± 45 minutes for NL4.3(iGFP)(ΔENV)(YP^-^) in HeLa cells. Recent studies have shown recruitment of ESCRTs is followed by release of the virion within a 20 second time window from the last ESCRT recruitment ^26^. In our experiments we suggest that release will only happen at the last recruitment of ESCRTs during the stuttering events on individual virions.

The recruitment of ALIX, CHMP4b and VPS4 with almost the same number of molecules under various late domain mutations argues that recruitment of ESCRTs is driven by a cooperative network which can be triggered through multiple entry points with identical net resulting recruitment. More recently binding of ubiquitin to Gag through ubiquitin ligases as well as membrane curvature where shown to play a role in recruitment of ESCRTs ^27-32^, however how these events are choreographed on the plasma membrane remains to be explored.

The stuttering recruitment of the full ESCRT machinery on the YP^-^ VLPs suggests that ALIX plays a major role during end stages of the ESCRT function and its function requires proper connection with the HIV Gag. Failure to connect properly with Gag results in disassembly and re assembly of the full ESCRT machinery. Therefore we argue that the prevalent linear biochemical interaction map between ESCRTs may unnaturally simplify the *in vivo* function of these interactions. There are up to five different ALIX interactions functioning at late stages of VLP assembly: i) A direct ALIX-Gag interaction through the YPXL late domain motif on p6 Gag ^2, 7, 15^, ii) A direct ALIX-Gag interaction through a binding site on NC Gag ^33,34,35^, iii) ALIX interactions with ubiquitin ^36,37^, iv) ALIX-TSG101 interactions ^3,4,38^, and v) interactions with ALIX itself, including relief of ALIX autoinhibition ^39,40^,opening of the V domain ^39^, and possibly ALIX dimerization ^41^. Our results suggest that the exact choreography of these interactions and what role they play during the function of the full ESCRT machinary can not simply be recruitment and remains to be visualized in vivo.

The ALIX homologue Bro1 in yeast is proposed to be recruited through interactions with Snf7 a yeast homologue of CHMP4 ^19,42^. ALIX has a Bro1 domain analogues to the yeast Bro1, along with a V domain and a PRR which does not exist in the yeast homologue Bro1 ^2,19^. The binding of late domain YPXL has been mapped to the ALIX V domain ^2^ and Cepp55 binds the PRR ^8,9^. Such apparent diversity had suggested that the recruitment and possibly function of ALIX is evolutionary separate from the yeast homologue Bro1. In contrast to this view, our observations showing that the late domain does not play a role in recruitment of ALIX is more in agreement with the findings in yeast where Bro1 was shown to regulate the function of ESCRT-III protein Snf7 during membrane scission^43^. While ALIX has been shown to be important in function of ESCRTs in all membrane scission reactions, a unified understanding of its function has been lacking. Based on all above data and available literature we suggest that ALIX plays a critical role during the final stages of membrane fission along with ESCRT-III and VPS4 proteins.

Live imaging of HIV as well as MVB budding has been previously used for visualizing recruitment of ESCRTs during membrane session events ^14,17,20,26,44-47^. Our study shows how disturbing previously characterized biochemical interactions can result in surprising recruitment profiles of ESCRTs observed in live cells and therefore underscores the usefulness of the imaging methods for further characterizing these interactions in vivo.

## Methods

### Cell Culture and Transfection

HEK 293T, HeLa and U2OS cells were maintained in Dulbecco’s modified Eagle’s medium (DMEM; Invitrogen, Carlsbad, CA) supplemented with fetal calf serum (10%), sodium pyruvate (1 mM) and L-glutamine (2 mM). HeLa cell lines stably expressing eGFP-tagged ALIX was maintained in the same medium supplemented with Genetecin (0.5 mg/mL) for selection. For TIRF experiments, cells were incubated in CO2-independent medium (LifeTechnologies).

Cells were seeded 18 h before transfection on sterile 4 chamber dishes at 60% confluency.Transfection was carried out using Lipofectamine2000 (Invitrogen) and DNA plasmid at the ratio of 3:1 in HeLa and 2:1 for 293T cells and U2OS cells with total DNA of 1500 ng for imaging and 2000 ng for western blot. For stable cell line, the sample was supplemented in CO2-independent medium and moved to the microscope for imaging 4h after transfection. In the case of transient co-transfection of DNA plasmids in normal HeLa, cells were used for imaging 6-7 hours after transfection. The cells were kept at 37°C during the imaging.

### siRNA transfections

HeLa cells were seeded at 40% confluency and were transfected 24 hours later with siRNA targeting Luciferase (CUGCCUGCGUGAGAUUCUCdTdT) or Alix (GAAGGAUGCUUUCGAUAAAUU) using Lipofectamine-2000. After 72 hours, cells were re-transfected with siRNA. Again after another 48 hours cells were transfected with siRNA along with the desired plasmid. Cells were imaged 7 hours later.

### Microscopy

Live images were acquired using iMIC Digital Microscope made by TILL photonics controlled by TILL’s Live Acquisition imaging software as previously described^17^. Two wavelengths of laser, 488 nm diode laser (Coherent, Saphire 488) and 561 nm diode-pumped solid state (DPSS) laser (Cobolt Jive, 561 nm Jive High Power), were used to excite eGFP and mCherry, respectively. Laser beams passed through an AOTF (acousto-optical tunable filter) and focused into a fiber which delivers the light to TILL Yanus digital scan head and then Polytrope II optical mode switch. Polytrope hosts a quadrant photodiode used for TIRF penetration depth calibration, which was set to 150 nm for the experiments in this manuscript. Once the penetration depths for the experiments are set at the beginning of acquisition, a feedback loop keeps the focus of the objective on the sample by constantly monitoring the position of the back reflected beam with respect to the original beam. We also rotated the TIRF illumination on the objective back focal plane 1 turn/exposure (TIRF360) to maximize homogeneity of the TIRF images.

### Microscopy data analysis

Images from the microscope were stored as TIFF files and analyzed using Matlab software (Mathworks) as described previously^17^. Intensity of the fluorescent signal collected from each diffraction limited spot is proportional to the number of molecules within that position, however the intensity is also proportional to the laser intensity, position of molecules with respect to glass during TIRF and substitution level of WT versus fluorescent molecules in each particular cell. To compare intensities of the ESCRT recruitments in between various cells and experimental conditions, average intensity of the HIV unaffiliated ESCRT recruitments at the plasma membrane were used to normalize the fluorescent intensities in between cells.

### Cell detachment experiments

U2OS cells were transfected with Gag-eGFP or Gag-eGFP(YP^-^) and observed by TIRF imaging. At 12 hours post-transfection, cells with VLPs were first imaged using TIRF and then cells were gently detached using TryplE (LifeTechnologies). Detachment was achieved by removing the medium and washing once with PBS; a thin layer of TryplE was added to cover cells to allow cell to detach. After few minutes, the glass was again imaged with released VLPs left on the glass support.

### Western blot analysis

Virion and cell lysates were separated on 4-15% acrylamide gels and transferred to nitrocellulose membranes. The blots were probed with anti-p24 (183-H12-5C, NIH AIDS Reagent Program), anti-eGFP (Santa Cruz), and infrared dye coupled secondary antibodies (LI-COR) were used for immunoprobing. Scanning was performed with the Odyssey infrared imaging system (LI-COR) in accordance with the manufacturer’s instructions at 700 or 800 nm, accordingly.

### Infectivity Assay

HEK 293T cells (60% confluent in 4 cm plates) were transfected using lipofectamine-2000 transfection with NL4.3 alone or along with ΔCMV-eGFP-flex-CHMP4B plasmid. The supernatant was harvested 48 hours later. Infectivity was measured by adding the supernatant to TZM-B1 cells (80% confluent). 48 hours later cells were lysed using britelite plus Reporter Gene Assay (Perkin Elmer) and luminosity was measured using a Cytation 5 microscope, experiments were carried out in triplicate.

### Statistics

All conditions tested in the manuscript contain >20 analyzed virus like particles in the supplementary figures and >40 virus like particles in the main figures. There was no data selection applied to the sample, therefore all data collected from the microscopy was analyzed and plotted in the figures.

### Availability of data

All data and reagents used in this study are available upon request, this includes the ΔCMV-eGFP-Flex-CHMP4B plasmid characterized in this study and its sequence which is available upon request and also includes the Matlab code used for analysis of the imaging data.

### Cell lines

The Hela, HEK 293T and U2OS cells were obtained from ITCC. TZM-b1 cells provided by NIH AIDS Reagent Program.

## Acknowledgements

This study was supported by NIH R01GM125444 to (SS).

## Author contributions

SG, MB and SS designed the experiments, SG performed experiments and analyzed data, MB created reagents, and SS wrote the manuscript.

## Materials & Correspondence

Requests should be sent to

## Supplementary Figures

**Figure S1:**
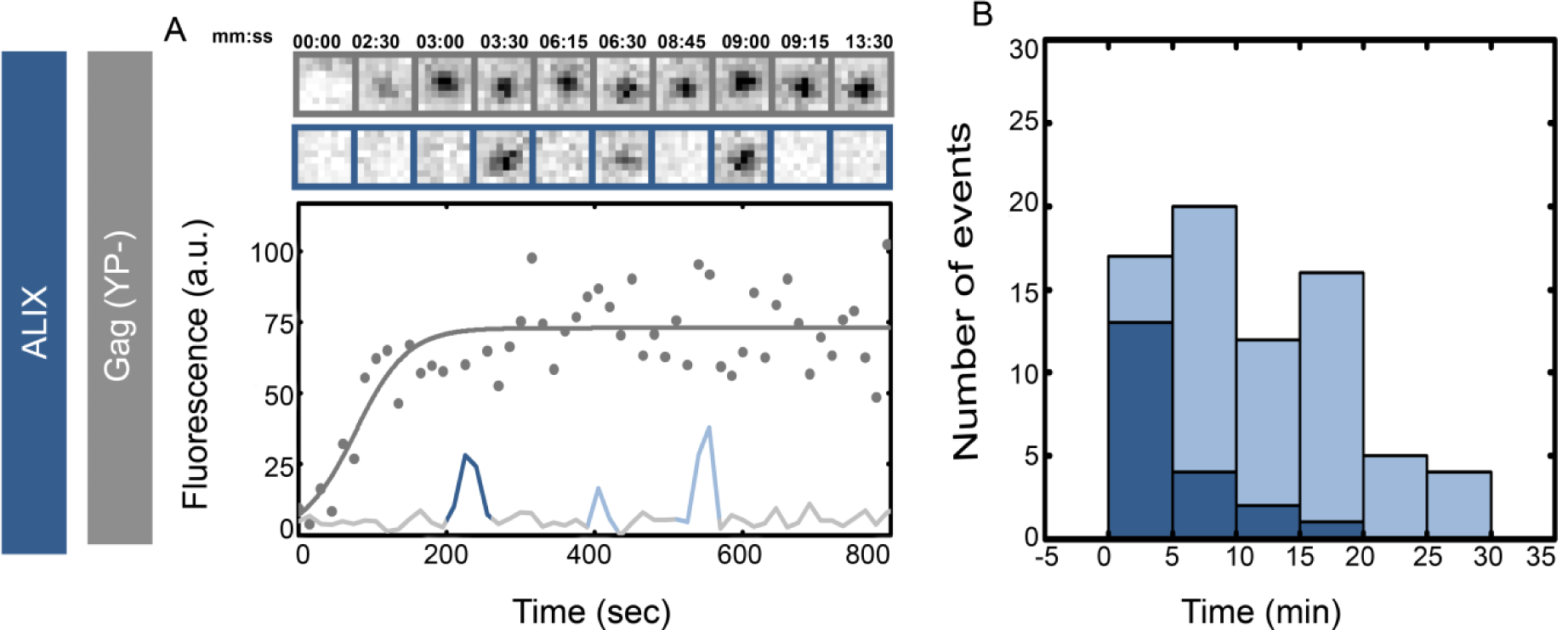
Figure1: Single versus multiple transient recruitments of ALIX into HIV Gag VLPs versus HIV Gag(YP^-^) VLPs. HeLa cells stably expressing ALIX-h30-eGFP were transfected with 300 ng of Gag(YP^-^) and 1200 ng of Gag(YP^-^)-mCherry and imaged 5 hours after transfection. (A) Shows assembly of individual representative VLPs with intensity plots and cropped TIRF images of the Gag (Top, Gray) and ALIX (bottom, Blue). Histograms of the first time (dark blue) and later recruitments (light blue) of ALIX are shown in (B). Majority of the first ALIX recruitment is within 1-10 minutes after the VLP assembly completes during YP^-^ assembly.

**Figure S2:**
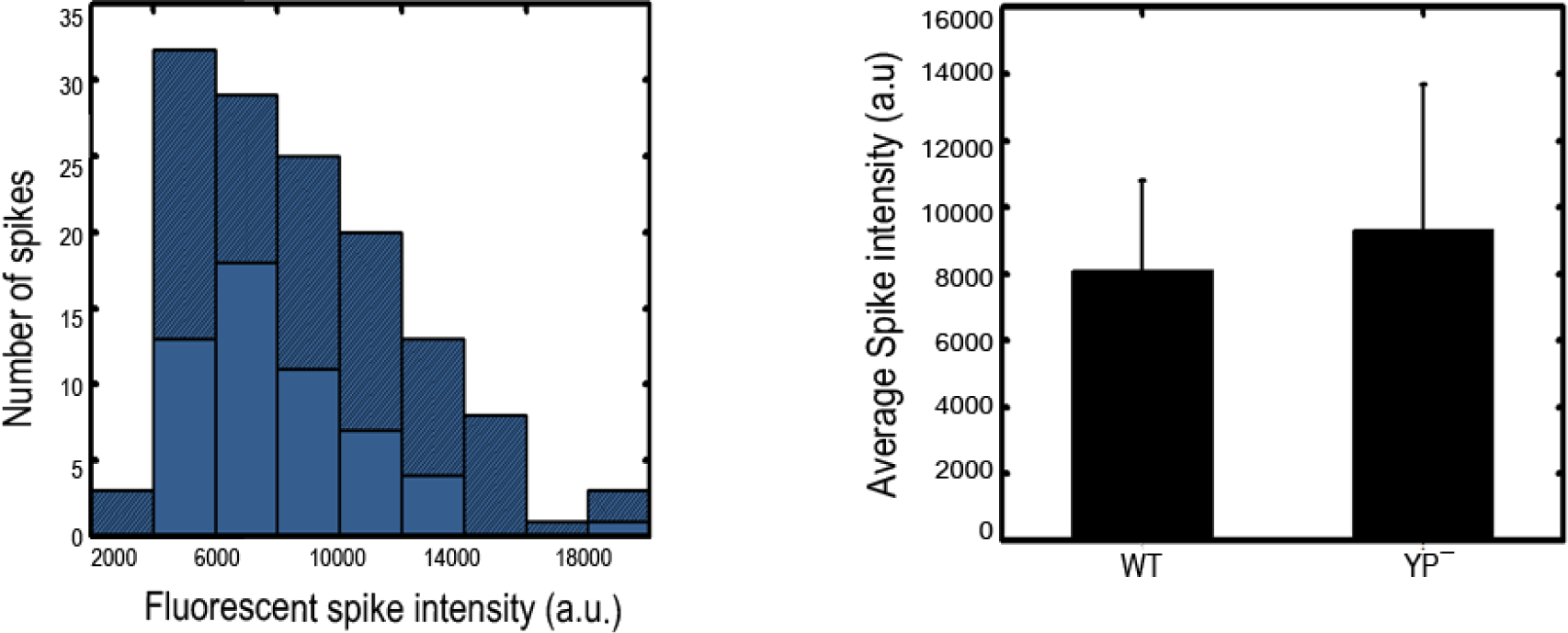
Distribution of the maximum fluorescence intensity of ALIX transient recruitment events. The maximum fluorescence intensity detected during recruitment is proportional to the number of molecules recruited to the site of assembly. Figures show a comparison between intensities during recruitment into HIV Gag-mCherry VLPs (light blue) versus HIV Gag(YP^-^)-mCherry VLPs (dark blue).

**Figure S3:**
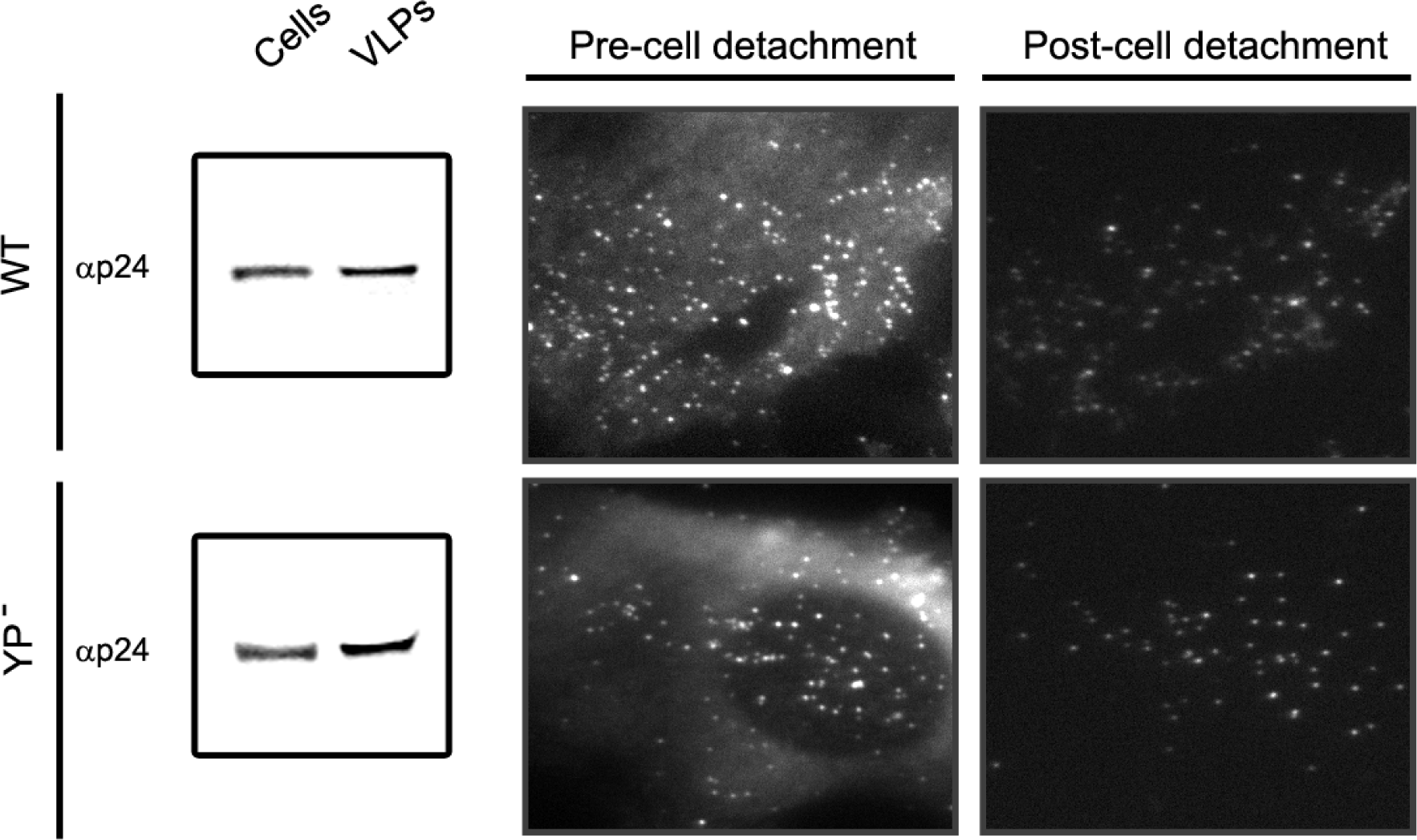
Similar release of Gag WT and (YP^-^) VLPs. Western blot conducted 24 hours after transfecting U2OS cells with Gag WT or (YP^-^) and immunoprobed with p24 (left panel) which shows no difference in release of VLPs from cells transfected with HIV Gag or HIV Gag(YP^-^). Single U2OS cells were imaged 12 hours post-transfection of HIV Gag-eGFP WT or HIV Gag(YP^-^)-eGFP using TIRF microscopy before and after cell detachment to visualize released VLPs (right panels)

**Figure S4:**
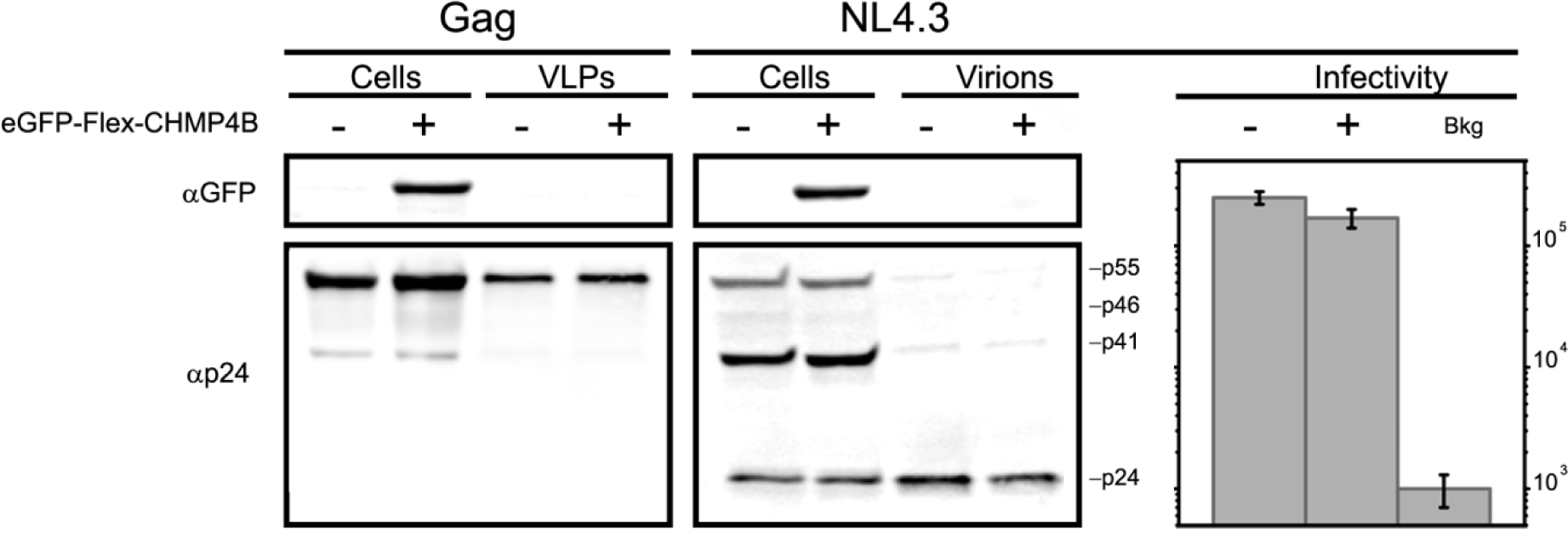
Release of HIV Gag VLPS or full HIV Virions with NL4.3 backbone was not affected on tagged CHMP4B co-expression. HEK 293T cells were transfected with ΔCMV-eGFP-flex-CHMP4b and HIV Gag or HIV NL4.3 respectively. Cells and supernatant were analyzed by western blots and probed with GFP and p24 (left panels). No difference is seen in release of HIV Gag VLPs while a slight decrease is detected in release of HIV NL4.3 virions from cells expressing eGFP-flex-CHMP4b. Infectivity assay for harvested NL4.3 virions using luciferase assay in TZM-b1 cells 48 hrs post infection (right panel). Bar graphs show a slight reduction of infectivity comparable to the reduction in virion release observed in western blot analysis.

**Figure S5:**
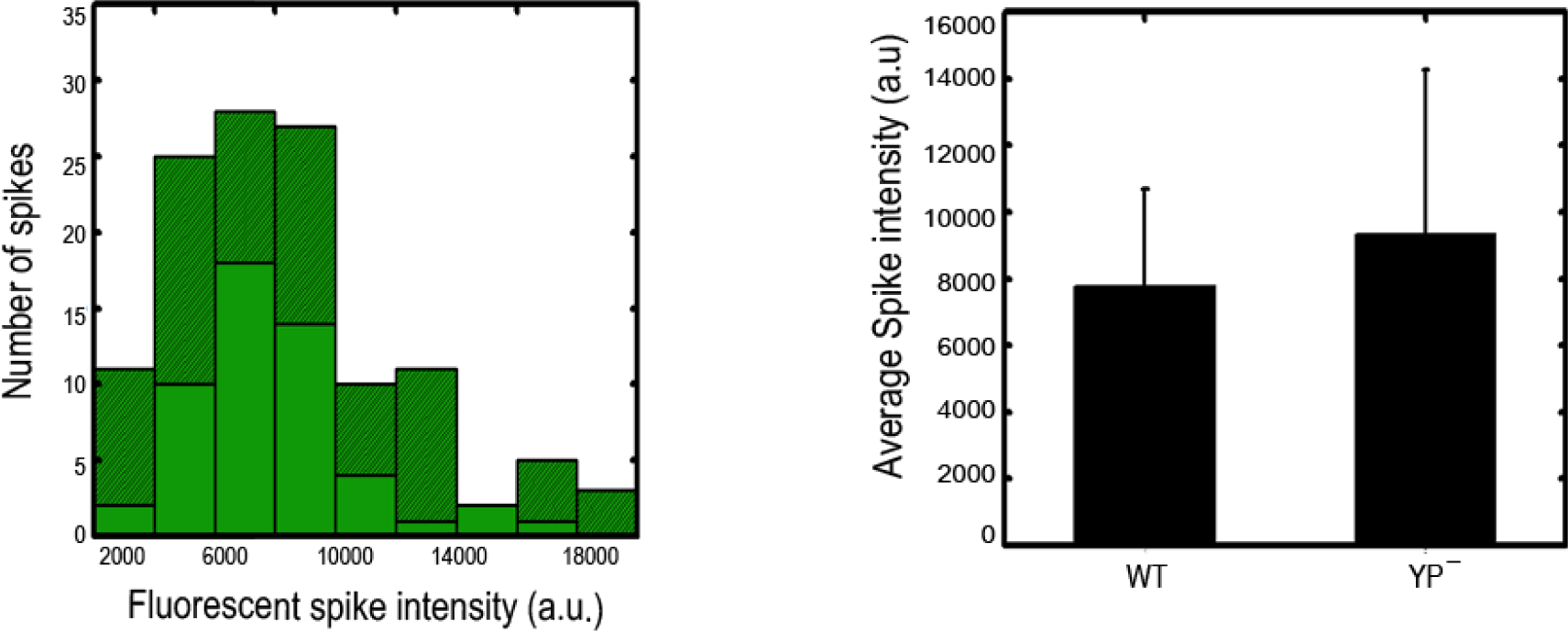
Distribution of the maximum fluorescence intensity of CHMP4 transient recruitment events. The maximum fluorescence intensity detected during recruitment is proportional to the number of molecules recruited to the site of assembly. Figures show a comparison between intensities during recruitment into HIV Gag-mCherry VLPs (light green) versus HIV Gag(YP^-^)-mCherry VLPs (dark green).

**Figure S6:**
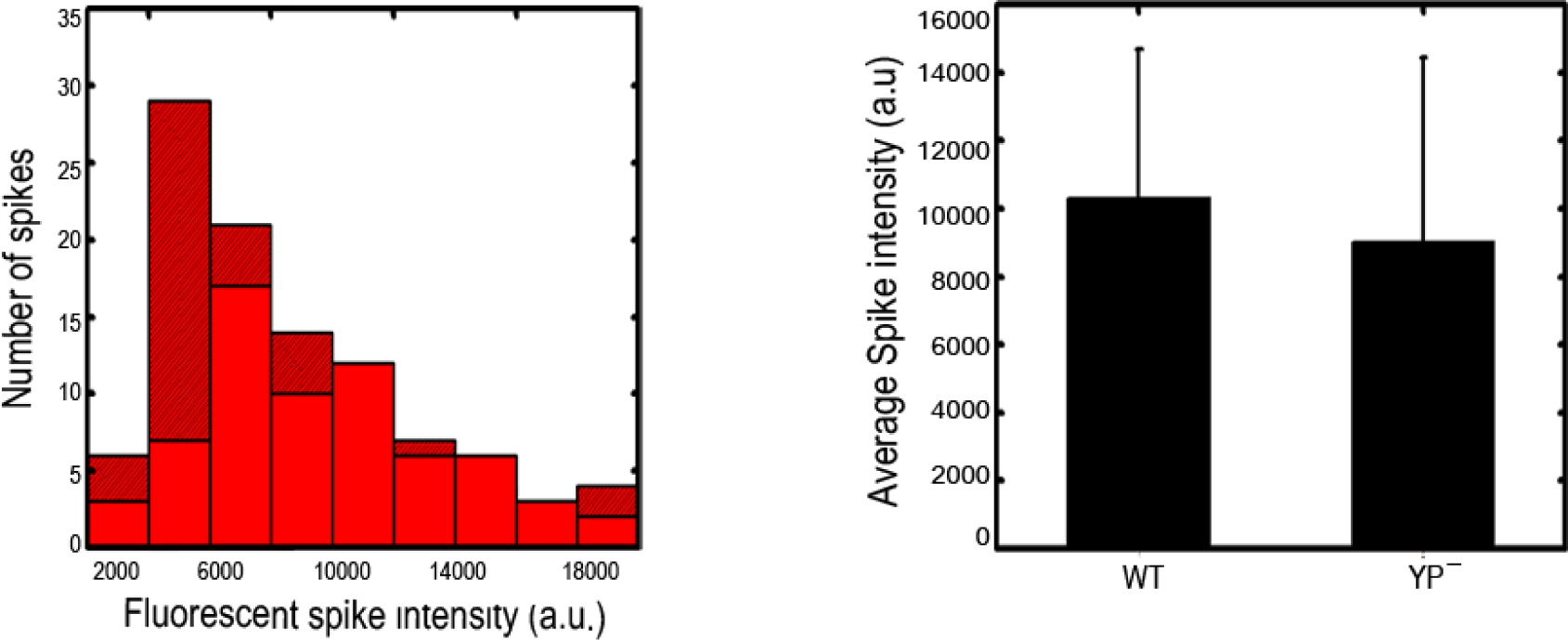
Distribution of the maximum fluorescence intensity of VPS4 transient recruitment events. The maximum fluorescence intensity detected during recruitment is proportional to the number of molecules recruited to the site of assembly. Figures show a comparison between intensities during recruitment into HIV Gag-mCherry VLPs (light red) versus HIV Gag(YP^-^)-mCherry VLPs (dark red).

**Figure S7:**
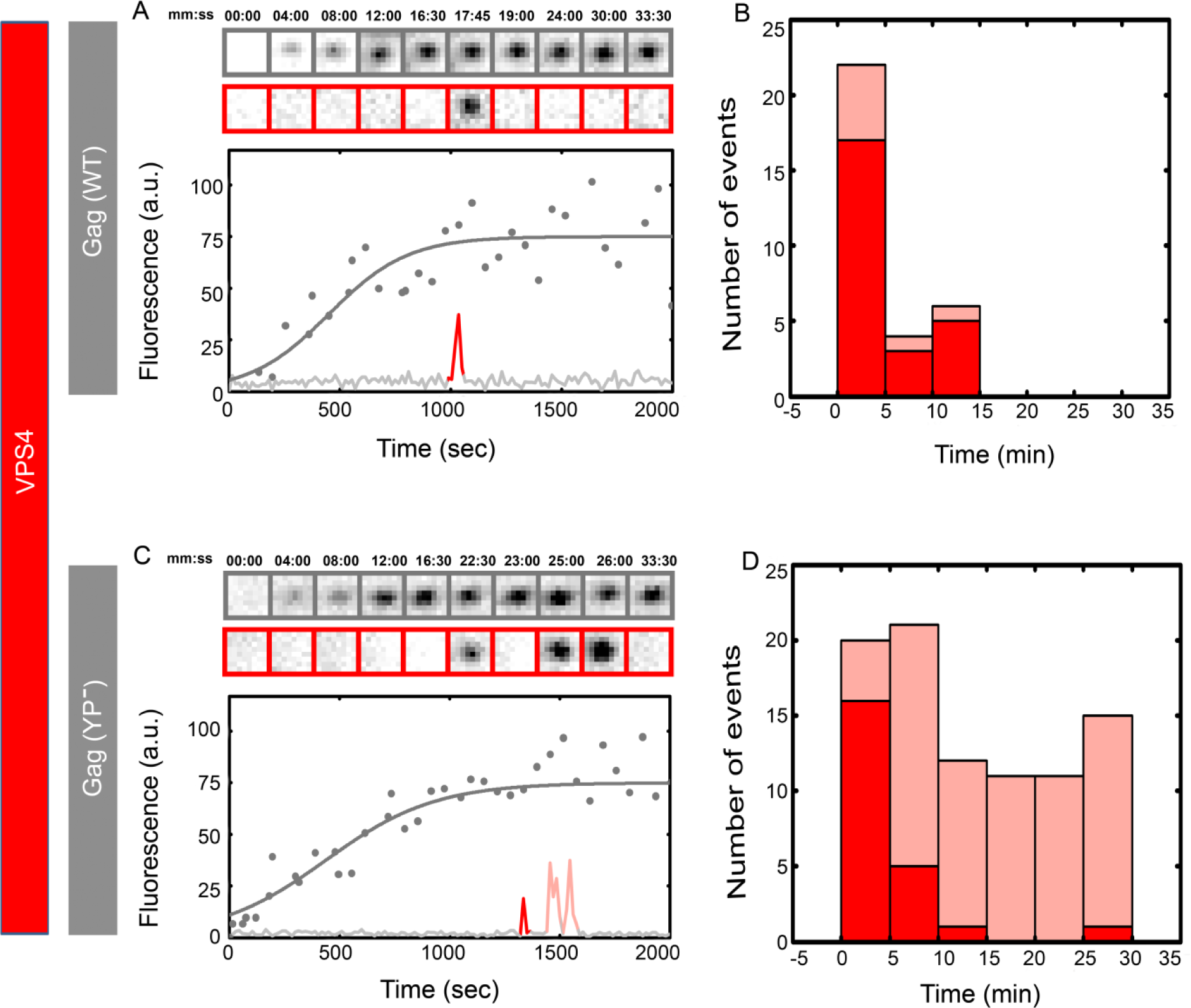
Recruitment of VPS4 into HIV Gag and HIV Gag(YP^-^) VLPs in U2OS cells. U2OS cells were transfected with 1200 ng of ΔCMV-VPS4-h37-mcherry and 300 ng of Gag-mCherry (A&B) and Gag(YP^-^)-mCherry (C&D) and imaged 7 hours after transfection. Assembly of individual representative VLPs are shown with intensity plots and cropped TIRF images of the Gag (Top, Gray) and VPS4 (bottom, Red) for (A) Gag-mChery VLPs and (C) Gag(YP^-^)-mCherry VLPs. Histograms of the first time (dark Red) and later recruitments (light Red) of VPS4 are shown for (B) Gag-mChery VLPs and (C) Gag(YP^-^)-mCherry VLPs. Majority of the first VPS4 recruitment is within 1-10 minutes after the VLP assembly completes during both YP^-^ and WT assembly.

**Figure S8:**
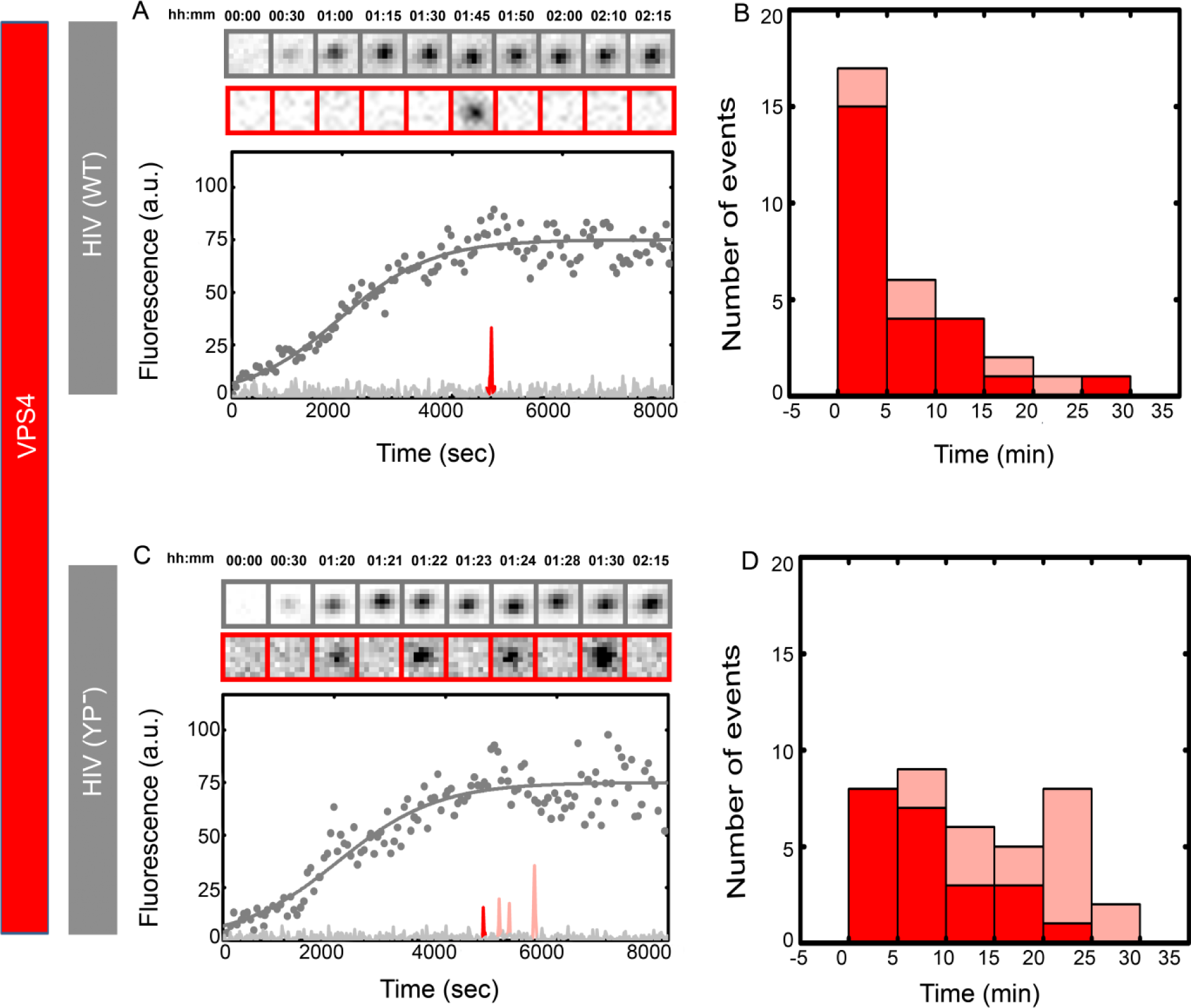
Transient recruitment of VPS4 into NL4.3(iGFP)(ΔENV) versus NL4.3(iGFP)(ΔENV) (YP^-^) VLPs. Hela cells were transfected with 300 ng of ΔCMV-VPS4-h37-mcherry and 1200 ng of NL4.3(iGFP)(ΔENV) (A&B) and NL4.3(iGFP)(YP^-^)(ΔENV) (C&D) and imaged 12 hours after transfection. Assembly of individual representative VLPs are shown with intensity plots and cropped TIRF images of the Gag (Top, Gray) and VPS4 (bottom, Red) for (A) NL4.3(iGFP)(ΔENV) VLPs and (C) NL4.3(iGFP)(YP^-^)(ΔENV) VLPs. Histograms of the first time (dark Red) and later recruitments (light Red) of VPS4 are shown for (B) NL4.3(iGFP)(ΔENV) VLPs and (C) NL4.3(iGFP)(YP^-^)(ΔENV). Majority of the first VPS4 recruitment is within 1-20 minutes after the VLP assembly completes during both YP^-^ and WT assembly.

**Figure S9:**
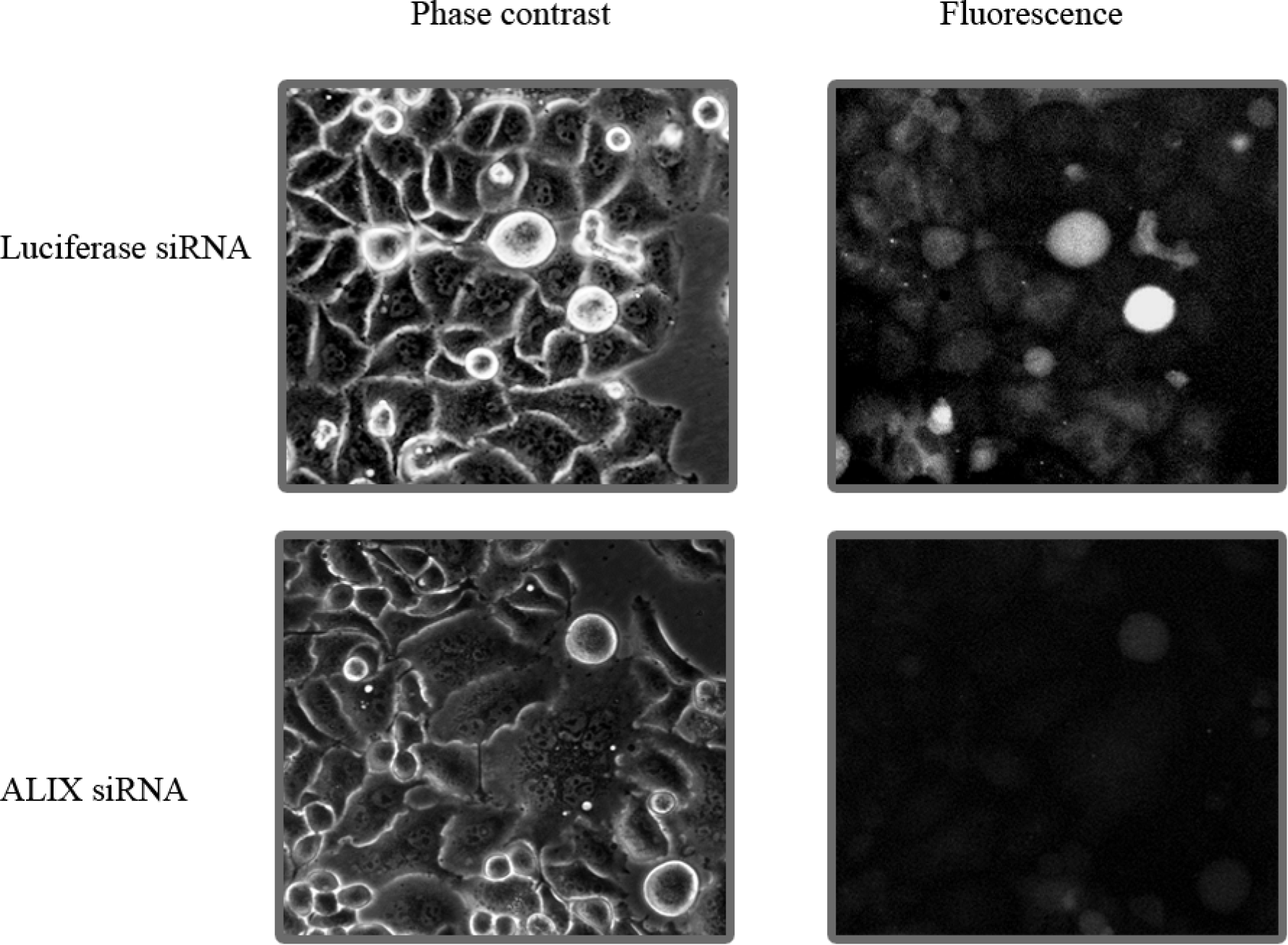
ALIX depletion using siRNA. Hela ALIX-h30-eGFP cells were treated with siRNA against ALIX or siRNA against luciferase(control) and imaged after 6 days. Fluorescence decrease and multinucleated cells are visualized in siRNA treatment against ALIX.

**Figure S10:**
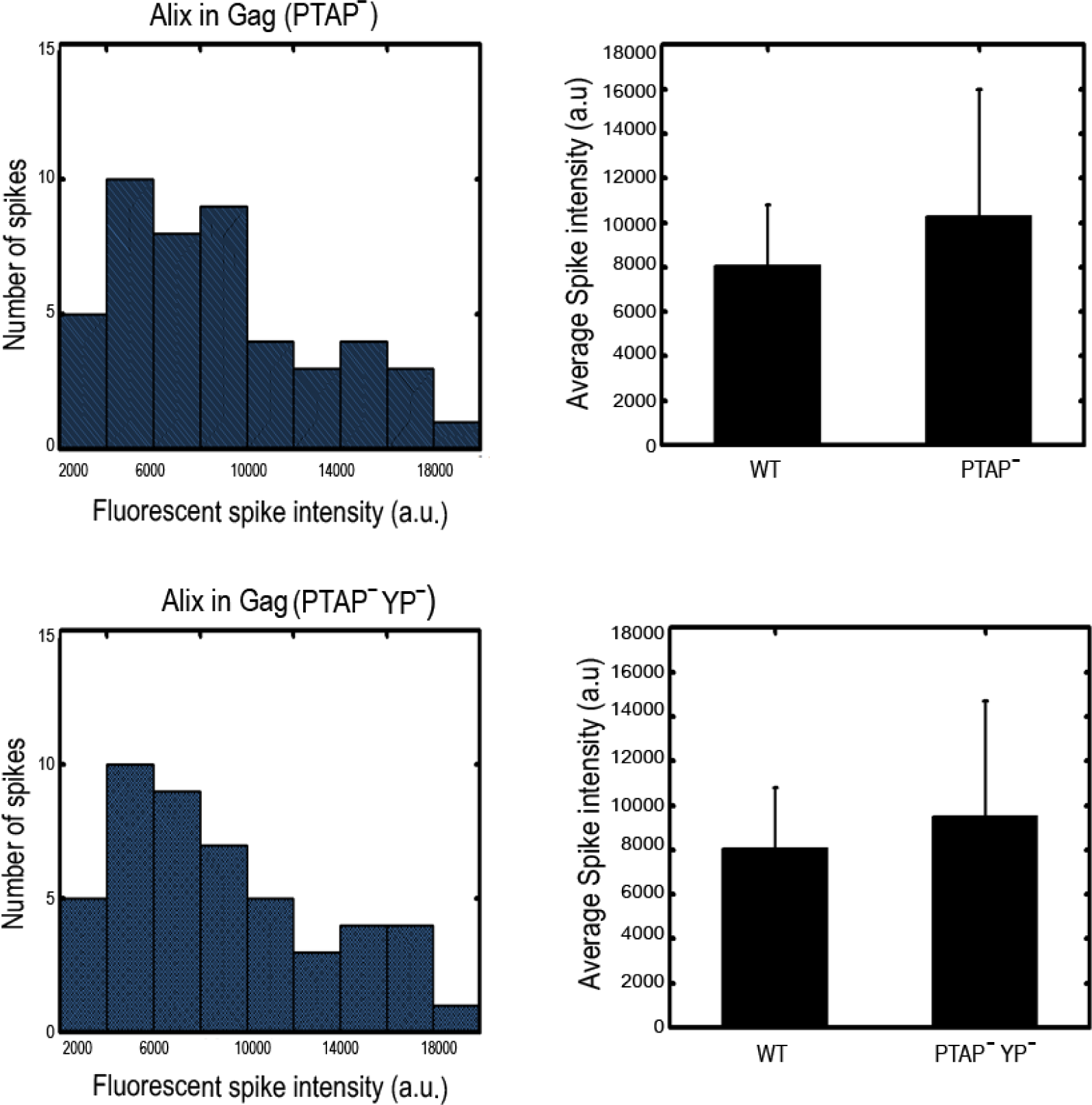
Distribution of the maximum fluorescence intensity of ALIX transient recruitment events into (PTAP^-^) and (PTAP^-^ + YP^-^) VLPs. The maximum fluorescence intensity detected during recruitment is proportional to the number of molecules recruited to the site of assembly.

## References

1. Henne, W. M., Stenmark, H. & Emr, S. D. Molecular Mechanisms of the Membrane Sculpting ESCRT Pathway. Cold Spring Harbor Perspectives in Biology 5(2013).

2. Fisher, R. D. et al. Structural and Biochemical Studies of ALIX/AIP1 and Its Role in Retrovirus Budding. Cell 128, 841–852 (2007).

3. Strack, B., Calistri, A., Craig, S., Popova, E. & Göttlinger, H. G. AIP1/ALIX Is a Binding Partner for HIV-1 p6 and EIAV p9 Functioning in Virus Budding. cell 114, 689–699 (2003).

4. Martin-Serrano, J., Yaravoy, A., Perez-Caballero, D. & Bieniasz, P. D. Divergent retroviral late-budding domains recruit vacuolar protein sorting factors by using alternative adaptor proteins. Proceedings of the National Academy of Sciences 100, 12414–12419 (2003).

5. Baietti, M. F. et al. Syndecan–syntenin–ALIX regulates the biogenesis of exosomes. Nature Cell Biology 14, 677 (2012).

6. Nabhan, J. F., Hu, R., Oh, R. S., Cohen, S. N. & Lu, Q. Formation and release of arrestin domain-containing protein 1-mediated microvesicles (ARMMs) at plasma membrane by recruitment of TSG101 protein. Proceedings of the National Academy of Sciences 109, 4146–4151 (2012).

7. Dores, M. R. et al. ALIX binds a YPX3L motif of the GPCR PAR1 and mediates ubiquitin-independent ESCRT-III/MVB sorting. The Journal of Cell Biology 197, 407–419 (2012).

8. Morita, E. et al. Human ESCRT and ALIX proteins interact with proteins of the midbody and function in cytokinesis. EMBO J 26, 4215–4227 (2007).

9. Carlton, J. G., Agromayor, M. & Martin-Serrano, J. Differential requirements for Alix and ESCRT-III in cytokinesis and HIV-1 release. Proceedings of the National Academy of Sciences 105, 10541– 10546 (2008).

10. Usami, Y., Popov, S. & Gottlinger, H. G. Potent Rescue of Human Immunodeficiency Virus Type 1 Late Domain Mutants by ALIX/AIP1 Depends on Its CHMP4 Binding Site. J. Virol. 81, 6614–6622 (2007).

11. McCullough, J., Fisher, R. D., Whitby, F. G., Sundquist, W. I. & Hill, C. P. ALIX-CHMP4 interactions in the human ESCRT pathway. Proceedings of the National Academy of Sciences 105, 7687–7691 (2008).

12. Scourfield, E. J. & Martin-Serrano, J. Growing functions of the ESCRT machinery in cell biology and viral replication. Biochemical Society Transactions 45, 613–634 (2017).

13. Votteler, J. & Sundquist, Wesley I. Virus Budding and the ESCRT Pathway. Cell Host & Microbe 14, 232–241 (2013).

14. Ku, P.-I., Bendjennat, M., Ballew, J., Landesman, M. B. & Saffarian, S. ALIX Is Recruited Temporarily into HIV-1 Budding Sites at the End of Gag Assembly. PLoS ONE 9, e96950 (2014).

15. Lee, S., Joshi, A., Nagashima, K., Freed, E. O. & Hurley, J. H. Structural basis for viral late-domain binding to Alix. Nat Struct Mol Biol 14, 194–199 (2007).

16. Jouvenet, N., Bieniasz, P. D. & Simon, S. M. Imaging the biogenesis of individual HIV-1 virions in live cells. Nature 454, 236–240 (2008).

17. Ku, P.-I. et al. Identification of Pauses during Formation of HIV-1 Virus Like Particles. Biophysical Journal 105, 2262–2272 (2013).

18. Ivanchenko, S. et al. Dynamics of HIV-1 Assembly and Release. PLoS Pathogens 5, e1000652 (2009).

19. Kim, J. et al. Structural Basis for Endosomal Targeting by the Bro1 Domain. Developmental Cell 8, 937–947 (2005).

20. Jouvenet, N., Zhadina, M., Bieniasz, P. D. & Simon, S. M. Dynamics of ESCRT protein recruitment during retroviral assembly. Nat Cell Biol 13, 394–401 (2011).

21. VerPlank, L. et al. Tsg101, a homologue of ubiquitin-conjugating (E2) enzymes, binds the L domain in HIV type 1 Pr55Gag. Proceedings of the National Academy of Sciences 98, 7724–7729 (2001).

22. Martin-Serrano, J., Zang, T. & Bieniasz, P. D. HIV-1 and Ebola virus encode small peptide motifs that recruit Tsg101 to sites of particle assembly to facilitate egress. Nat Med 7, 1313–1319 (2001).

23. Garrus, J. E. et al. Tsg101 and the Vacuolar Protein Sorting Pathway Are Essential for HIV-1 Budding. cell 107, 55–65 (2001).

24. Huang, M., Orenstein, J. M., Martin, M. A. & Freed, E. O. p6Gag is required for particle production from full-length human immunodeficiency virus type 1 molecular clones expressing protease. Journal of Virology 69, 6810–6818 (1995).

25. Bendjennat, M. & Saffarian, S. The Race against Protease Activation Defines the Role of ESCRTs in HIV Budding. PLoS Pathogens 12, e1005657, doi:10.1371/journal.ppat.1005657 (2016).

26. Johnson, D. S., Bleck, M. & Simon, S. M. Timing of ESCRT-III protein recruitment and membrane scission during HIV-1 assembly. eLife 7, e36221, doi:10.7554/eLife.36221 (2018).

27. Morita, E. et al. ESCRT-III Protein Requirements for HIV-1 Budding. Cell Host & Microbe 9, 235- 242, doi:DOI:10.1016/j.chom.2011.02.004 (2011).

28. Lee, I.-H., Kai, H., Carlson, L.-A., Groves, J. T. & Hurley, J. H. Negative membrane curvature catalyzes nucleation of endosomal sorting complex required for transport (ESCRT)-III assembly. Proceedings of the National Academy of Sciences 112, 15892-15897, doi:10.1073/pnas.1518765113 (2015).

29. Chung, H.-Y. et al. NEDD4L Overexpression Rescues the Release and Infectivity of Human Immunodeficiency Virus Type 1 Constructs Lacking PTAP and YPXL Late Domains. Journal of Virology 82, 4884–4897 (2008).

30. Mercenne, G., Alam, S. L., Arii, J., Lalonde, M. S. & Sundquist, W. I. Angiomotin functions in HIV-1 assembly and budding. eLife 10.7554, doi:10.7554/eLife.03778 (2015).

31. Sette, P., Jadwin, J. A., Dussupt, V., Bello, N. & Bouamr, F. The ESCRT-associated protein Alix recruits the ubiquitin ligase Nedd4-1 to facilitate HIV-1 release through the LYPXnL L domain motif. J. Virol., JVI.00634-00610, doi:10.1128/jvi.00634-10 (2010).

32. Weiss, E. R. et al. Rescue of HIV-1 Release by Targeting Widely Divergent NEDD4-Type Ubiquitin Ligases and Isolated Catalytic HECT Domains to Gag. PLoS Pathog 6, e1001107 (2010).

33. Sette, P., Dussupt, V. & Bouamr, F. Identification of the HIV-1 NC Binding Interface in Alix Bro1 Reveals a Role for RNA. Journal of Virology 86, 11608–11615 (2012).

34. Dussupt, V. et al. The Nucleocapsid Region of HIV-1 Gag Cooperates with the PTAP and LYPXnL Late Domains to Recruit the Cellular Machinery Necessary for Viral Budding. PLoS Pathog 5, e1000339 (2009).

35. Sette, P. et al. HIV-1 Nucleocapsid Mimics the Membrane Adaptor Syntenin PDZ to Gain Access to ESCRTs and Promote Virus Budding. Cell Host & Microbe 19, 336–348 (2016).

36. Dowlatshahi, Dara P. et al. ALIX Is a Lys63-Specific Polyubiquitin Binding Protein that Functions in Retrovirus Budding. Developmental Cell 23, 1247–1254 (2012).

37. Keren-Kaplan, T. et al. Structure-based in silico identification of ubiquitin-binding domains provides insights into the ALIX-V:ubiquitin complex and retrovirus budding. EMBO J 32, 538–551 (2013).

38. von Schwedler, U. K. et al. The Protein Network of HIV Budding. cell 114, 701–713 (2003).

39. Zhai, Q. et al. Activation of the Retroviral Budding Factor ALIX. J. Virol., JVI.02653-02610 (2011).

40. Zhou, X. et al. The CHMP4b- and Src-docking sites in the Bro1 domain are autoinhibited in the native state of Alix. Biochem J 418, 277–284 (2009).

41. Pires, R. et al. A Crescent-Shaped ALIX Dimer Targets ESCRT-III CHMP4 Filaments. Structure 17, 843–856 (2009).

42. Odorizzi, G., Katzmann, D. J., Babst, M., Audhya, A. & Emr, S. D. Bro1 is an endosome-associated protein that functions in the MVB pathway in Saccharomyces cerevisiae. Journal of Cell Science 116, 1893–1903 (2003).

43. Wemmer, M. et al. Bro1 binding to Snf7 regulates ESCRT-III membrane scission activity in yeast. The Journal of Cell Biology 192, 295–306 (2011).

44. Bleck, M. et al. Temporal and spatial organization of ESCRT protein recruitment during HIV-1 budding. Proceedings of the National Academy of Sciences 111, 12211–12216 (2014).

45. Prescher, J. et al. Super-Resolution Imaging of ESCRT-Proteins at HIV-1 Assembly Sites. PLoS Pathog 11, e1004677 (2015).

46. Baumgartel, V. et al. Live-cell visualization of dynamics of HIV budding site interactions with an ESCRT component. Nat Cell Biol 13, 469–474 (2011).

47. Adell, M. A. Y. et al. Recruitment dynamics of ESCRT-III and Vps4 to endosomes and implications for reverse membrane budding. eLife 6, e31652, doi:10.7554/eLife.31652 (2017).

